# Sleep-dependent modulation of thalamic brain temperature estimates in humans

**DOI:** 10.64898/2026.10.08.757550

**Authors:** Marieke MB Hoekstra, Mick Lehmann, Dominic Gascho, Patrik Verstreken, Christina Schmidt, Hans-Peter Landolt, Niklaus Zölch

## Abstract

Mammalian thermal physiology is geared towards dissipating heat during sleep. Brain temperature typically decreases during non-rapid-eye-movement (NREM) sleep in all mammalian species measured to date, yet non-invasive confirmation in healthy humans is lacking: the only available human study used invasive recordings in epileptic patients without a wake control condition. Magnetic resonance spectroscopy (MRS) enables brain temperature mapping non-invasively, using the temperature-sensitive chemical shift of water relative to N-Acetyl-Aspartate as an internal reference. First, we validated that with metabolite cycling (MC), a MRS approach that enables the simultaneous acquisition of metabolite and water information without conventional water suppression, can be used to determine estimates of brain temperature in an *ex vivo* sheep head. Next, we determined that by using a temporal resolution of ∼4 minutes, we obtained high-quality spectra without losing excessive temporal resolution. We then re-analyzed an existing set of MRS data (acquired with the MC sequence) of the thalamus in young adults (n=9) who slept in the evening for up to 2 hours in the scanner, to assess sleep-evoked changes in estimates of brain temperature. Of the nine participants, five participants took part in the subsequent wake-only experiment, allowing to control for the potential impact of circadian time and body posture on estimates of brain temperature. Sleep-wake states were quantified with standard polysomnography, confirming that the participants managed to sleep in the scanner. On average, they spent 39.4±15.7 min in N2 sleep, 24.8±18.1 min in N3 sleep, 10.0±4.9 min in N1 sleep, and 37.0±19.6 min awake. Estimates of brain temperature were correlated across the sleep and wake-only experiment within the same participant (R^2^=0.29, P<0.0001). When participants entered NREM sleep, estimates of brain temperature decreased by 0.2°C. Moreover, the deeper NREM sleep, the more pronounced the decrease in estimates of brain temperature (P=0.014). These data suggest that an evening nap consisting of NREM sleep decreases thalamic temperature. Considering the profound impact temperature can have on brain functioning, these brain temperature changes could be a pathway through which sleep impacts the brain.

## Introduction

The regulation of sleep and temperature are intricately linked. Sleep preferentially occurs when body temperature is on its steepest nightly decrease ^1^, and the higher core body temperature at sleep onset, the longer sleep duration ^2^. Peripheral vasodilation, a thermoregulatory process that is essential to promote heat loss from the core, is a characteristic of sleep initiation in mammals^3^, and impaired peripheral vasodilation is associated with an increased sleep onset latency^4^, which can be partially resolved by warming extremities (e.g. wearing socks in bed) ^5^. Along these lines, individuals with poor sleep quality have elevated core body temperature prior and during sleep compared to participants with good sleep quality ^6^. Thus, there is a tight link between peripheral thermoregulation and sleep initiation and continuation.

Beyond peripheral temperature, the effect of sleep on brain temperature has been studied extensively using invasive thermosensitive probes in a plethora of mammalian species. These studies showed that there is a remarkable close and fast association between brain temperature and sleep. Specifically, in mice brain temperature decreases during sleep, albeit with opposing dynamics across non-rapid-eye-movement (NREM) and rapid-eye-movement (REM) sleep: it falls during non-rapid eye movement (NREM) sleep (in 90s approximately -0.2°C) and rises during REM sleep (in 90s +0.4°C). Because NREM sleep is the dominant sleep state in mammalian species, this state specific evoked changes result in a net sleep-induced decrease of 1.5-2°C across the day ^7^. In mice, these changes in brain temperature are so much dependent on sleep-wake state, that we can predict sleep-wake states with 91% accuracy based on 4-sec epochs of brain temperature ^8^. Brain temperature can also track specific neuronal states, such as sleep spindles during NREM sleep and high theta and gamma power in wakefulness ^9^. Suggesting a functional role for sleep-dependent changes in brain temperature comes from elegant work of the Buszáki lab: modulation of hippocampal temperature results in altered number of sharp-wave ripples, a neuronal signature of information transfer during sleep ^10^.

Despite the wealth of literature pointing to a close association between temperature and sleep at different levels and across various mammalian species, specific data on the impact of sleep on human brain temperature is surprisingly sparse. To date, there is only one study that addressed this topic by inspecting brain temperature changes in epilepsy patients with thermal probes placed intracranially in the para-hippocampal gyrus while simultaneously recording sleep with polysomnography. In this study, brain temperature decreased during sleep time, resulting in a daily amplitude of >0.5°C ^11^. Whether these results hold when controlling for body posture, such as is done in a constant routine protocol, and sleep-wake state, with a circadian time matched wake control, is not clear.

The sparsity in studies is mainly because non-invasive measurement of brain temperature in humans has been challenging. More specifically, direct measurements are restricted to invasive thermal probes such as the one used in the study above, limiting their application to patients. Proton (¹H) magnetic resonance spectroscopy (MRS) offers an elegant solution to this limitation, exploiting the well-established temperature dependence of the water proton resonance frequency (∼0.01 ppm/°C) relative to temperature-stable metabolite references such as N-acetyl-aspartate (NAA) whose chemical shift is independent of temperature ^12–15^. Zhu and colleagues improved MRS-based temperature estimation (MRS thermometry) using an ultra-high-field 11-T magnetic resonance (MR) scanner in the rat brain, achieving an estimated accuracy of approximately 0.1°C ^16^. Since then, this technique was applied in a range of clinical and research contexts, including stroke, hypothermia monitoring, and neurodevelopmental studies ^17^. While a recent study confirmed that daily oscillations in brain temperature during wakefulness can be detected with ¹H-MRS ^18^, it remains unclear whether and to what extent these daily temperature changes can be explained by sleep-wake states. We hypothesized that thalamic temperature estimates would decrease during NREM sleep compared with time-matched wakefulness. To test this hypothesis, we analyzed a previously published dataset in which sleep-wake state was co-detected with MRS spectra in the thalamus ^19^, a brain region important for sleep induction and maintenance ^20^.

## Material and methods

### Participants and experimental procedure

A detailed description of the participants and experimental procedure can be found in Lehman et al. 2019 ^19^. In brief, fifteen participants were recruited of which six participants (2 unable to sleep, 4 slept <15 min) were excluded, yielding n = 9 (5F/4M, mean age 22.3±2.2 years). Participants sleeping >15 min in N2/N3 underwent a second, wake-only control session (see Figure 1). One week before the recording night, participants maintained habitual sleep-wake schedules, verified by actigraphy and sleep logs, with sleep duration reduced by 2 h on the final night to elevate sleep pressure. On the experimental night, polysomnography (32-channel EEG, EOG, EMG, ECG) was applied ∼3 h before habitual bedtime; structural MR imaging began ∼2 h before bedtime. Participants stayed awake for ≥20 min during initial MRS acquisition, then slept; MRS/PSG continued until movement precluded further scanning. The Ethics Committee of the Canton of Zurich approved the study. All participants gave written informed consent before screening.

**Figure 1.**
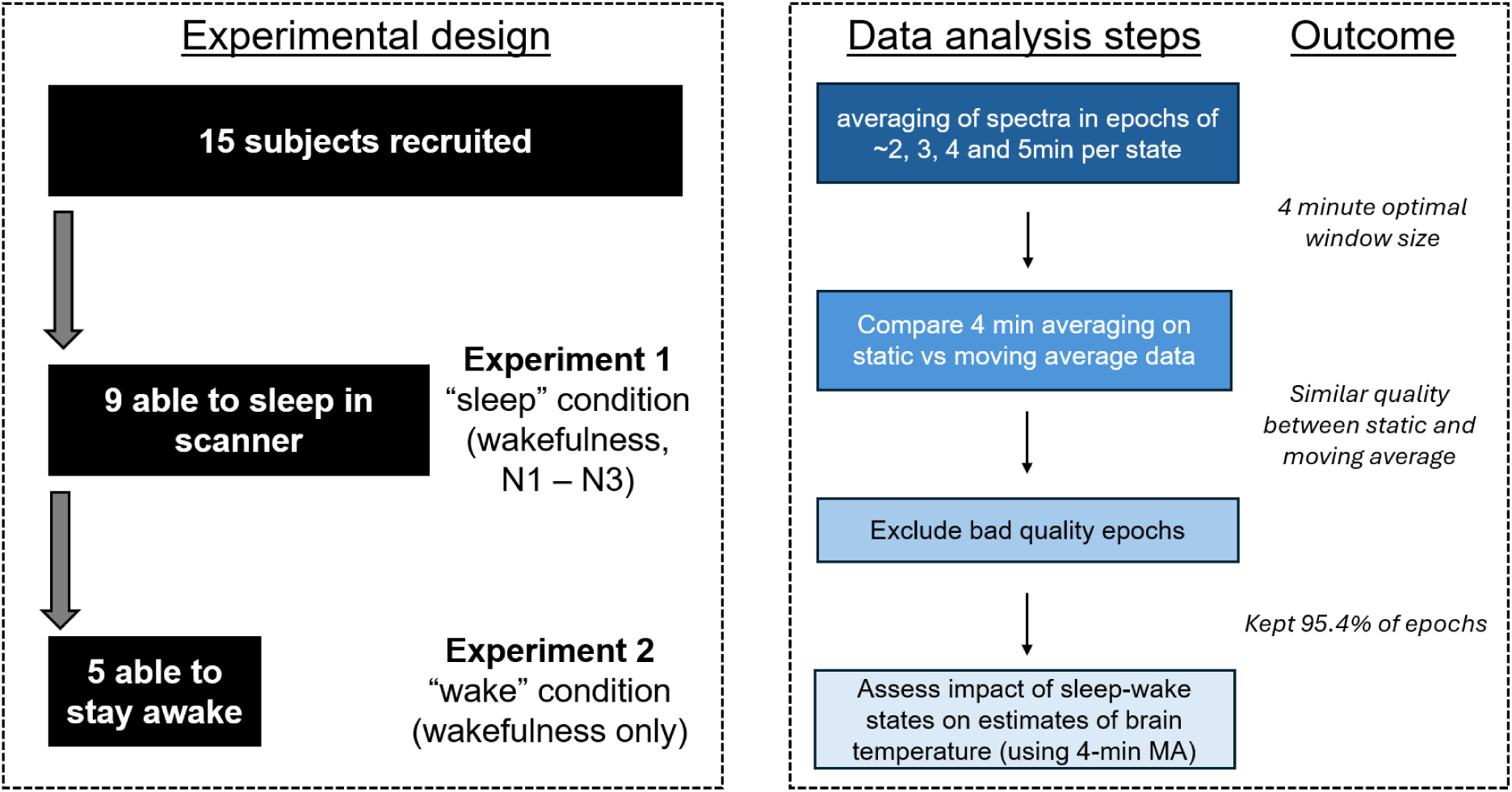
Experimental design and analysis. Left panel: overview of the experimental design of the study; right panel: overview of the sequential analyses steps.

### Polysomnographic Recording and Artifact Correction

A detailed description of EEG acquisition can be found in Lehman et al. 2019 ^19^. Briefly, wakefulness and sleep were confirmed by MR-compatible polysomnography (BrainAmp, BrainCap, BrainAmp ExG MR; Brain Products GmbH, Gilching, Germany; impedances <10 kΩ), synchronized to the scanner clock, sampled at 5 kHz (0.01-200 Hz), referenced to Cz. Offline correction (BrainVision Analyzer 2.0) applied average artifact subtraction for MRS artifacts (21-interval sliding average) and cardioballistogram template correction (21-cycle moving average, ECG-derived) ^21^, both pre-implemented in the software. Data were filtered (0.3-35 Hz, 48 dB/octave), down-sampled to 512 Hz, and scored offline in 30-s windows by two independent raters per standard criteria ^22^. Sleep onset was defined by finding the first sleep occurrence (in any state), followed by uninterrupted N2 sleep for at least 6 minutes (min). Due to the MR-induced artefacts on the EEG traces, reliable spectral analysis on brain activity was not possible.

### MRS acquisition

*Scanner specs* ^1^H-MRS measurements were performed on a Philips Achieva 3T whole-body scanner equipped with a 32-channel receive-only phased-array head coil (Philips Healthcare, Best, The Netherlands). Non-water suppressed, PRESS-localized ^1^H-MRS spectra (2500 ms repetition time, 32 ms echo time and 2000 Hz spectral bandwidth) in the thalamus were obtained using the metabolite cycling (MC) technique ^23^, optimized with six very selective saturation (VSS) pulses to minimize errors in chemical-shift displacement and to achieve consistent localization volumes across all metabolites of interest. The nominal voxel-of-interest (VOI) size was 22 × 13 × 16 mm^3^. Accounting for the VSS pulses, the resulting VOI size was 14.4 × 10.0 × 10.4 mm^3^ (≈ 1.5 mL).

In the MC technique, spectral regions on either side of the water resonance are alternately inverted in consecutive transients located upfield and downfield of the water resonance, are selectively inverted in consecutive transients, while the water signal remains largely unaffected. The metabolite spectrum is subsequently obtained by subtracting transients acquired with opposite inversion profiles, thereby isolating metabolite signals without the need for water suppression.

Because the water resonance remains unchanged between transients, it can be used to correct for frequency and phase variations occurring between consecutive transients. Furthermore, the continuously present water signal enables monitoring of changes in the water resonance frequency and, in principle, allows temperature changes to be tracked with a temporal resolution corresponding to the repetition time (TR). Absolute temperature estimation, however, requires determination of the chemical shift difference between water and NAA. Due to its substantially lower concentration than water, the NAA resonance exhibits a lower signal-to-noise ratio, requiring averaging of multiple transients for reliable determination of its chemical shift. Consequently, the temporal resolution of the temperature measurement is determined by both the number of averaged transients (hereafter referred to as epochs) and the TR. Throughout this manuscript, averaged sets of transients are referred to as epochs. The minimum number of transients required for reliable temperature estimation was determined experimentally (see Results, section 2: Trading spectral quality for temporal resolution).

Despite the potential advantages of having access to both the resonance frequency of water and NAA, making MC suitable for temperature measurements, this technique has not previously been applied to MRS thermometry to our knowledge. Therefore, the MC-based temperature measurement and the associated analysis methodology were first evaluated in sheep heads.

### Sheep experiment

We first assessed whether our specific acquisition and analysis pipeline would allow us to quantify brain temperature changes using MRS. To this end, we conducted absolute temperature measurements in post-mortem sheep heads simultaneously with MRS thermometry.

All specimens were collected directly after the end of another animal study which was performed according to the Swiss animal welfare act (animal license number ZH234/17). The sheep were deeply anesthetized with pentobarbital (Pentobarbitalum natricum, Esconarkon ad us. vet. [100–150 mg/kg bodyweight], Streuli Pharma AG). After confirmation of death by the absence of heartbeat and pupillary reflex, the head was severed, placed in a heat-insulating box and transported to the research facility to commence the MRS experiments.

To assess whether we could reliably predict proxies of brain temperature, we applied MRS with the MC PRESS sequence on severed sheep heads (n=3) and measured the temperature decreasing towards ambient temperature in brain tissue. MRS measurements started approximately 60 min after decapitation. A MRS voxel was placed on the level of the thalamus on the right hemisphere with a resulting VOI size of 12 × 13 × 12 mm^3^ (≈ 1.9 mL) accounting for VSS pulses and averaged 64 transients for one temperature estimate (temporal resolution: 2 min 26 sec64 * TR (2.288 s)). Contralateral of the VOI in which we applied MRS thermometry, a MR-suitable fiber optic temperature probe (GaAs-crystal, polyimide tip with 0.7-mm diameter, 2-m length, accuracy of relative temperature ±0.2 C; provided by Rugged Monitoring, Canada) was inserted, also in the thalamus. We analyzed only temperature values above >32°C as measured by the temperature probe, staying relatively close to physiological brain temperature values and thereby minimizing a potential influence of postmortem effects.

### Chemical Shift Determination using LCModel

To estimate temperature from the MRS spectra, we determined the chemical shift difference between the water and NAA resonances using LCModel, which estimates metabolite parameters by fitting the measured spectrum as a linear combination of reference metabolite spectra. A basis set containing 20 metabolites was simulated, using FID-A (lcm_gamma_std_press_thal see https://github.com/arcj-hub/BasisSetSimulation). The water - NAA chemical shift difference is not directly provided by LCModel, but can be determined from two frequency shift parameters reported in the LCModel output. In the first step of the analysis, the spectrum is shifted such that the major landmarks in the spectrum, including the NAA and creatine (Cr) resonances, are aligned with the expected chemical shifts. This shift is reported as “data shift” in the LCModel .table output file. The increment of this shift is determined by the frequency spacing between points in the spectrum, given by 1/Tacq, corresponding to ∼0.004 ppm in the present study. In the second step, LCModel optimizes the frequency shift of each metabolite on a much finer spectral resolution. The resulting frequency shift for NAA (“NAA shift”) is reported in the LCModel .print output file with a precision of 0.1 ppb. The water-NAA chemical shift difference (Δ), referred to as “Δ chemical shift”, was calculated from these two parameters, and subsequently converted to temperature estimates as described below.

### Conversion of MRS chemical shift values into an estimate of absolute brain temperature

To convert our measurements of from Δ chemical shift in the human brain MRS thermometry data to a proxy of brain temperature, we applied the following formula:

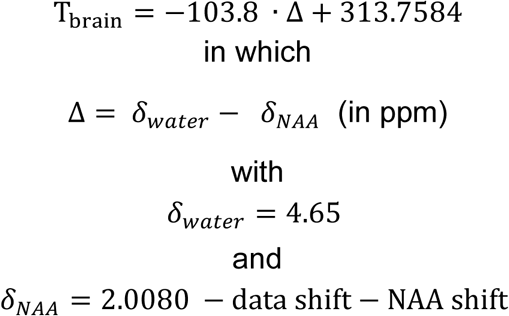

These constants, -103.8 and 313.7584, are based on the benchmarking study from Zhu et al. 2008, in which brain temperature in rats was estimated at high resolution using a 11.7T scanner, and compared to temperature measurements using an optical fiber ^16^. We decided to adopt Zhu’s coefficients for comparability with the human MRS thermometry literature, in line with a recent postmortem MRS thermometry study in which the Zhu calibration showed the best agreement with direct probe measurements among several published calibration approaches^15^.

### Trade-off between temporal resolution and spectral quality

To find the optimum between good quality data and high temporal resolution in the human sleep experiment, we extracted four quantitative measures of spectral quality: signal-to-noise ratio (SNR), linewidth, estimates of the relative Cramér-Rao Lower Bound (CRLB) and variation in brain temperature, referred to as “Δ temperature root mean square (RMS)”. The SNR was calculated as follows: SNR = max Amplitude [NAA] / SD of signal free range [0 - 0.2ppm]; linewidth is calculated as: full width at half maximum for the water peak. The CRLB is calculated for the NAA peak area and an output of LCModel, providing an estimate of the minimum achievable standard deviation of a modeled quantity across repeated measurements under identical experimental conditions^24^. Finally, we quantified within-participant variability of temperature estimates as the maximum absolute difference between consecutive epochs, and as the root mean square of successive epoch-to-epoch differences (Δ temperature RMS). Landolt et al. detected a group-based average daily amplitude of ∼0.5°C (with individuals exhibiting a daily amplitude of ∼1.5°C) ^11^. Hence, epoch-to-epoch differences of ≥0.5°C are unlikely reflect true biological measurements but could rather reflect a measurement error. Hence, an increased Δ temperature RMS indicates lower data quality. These four variables, SNR, linewidth, CRLB and Δ temperature RMS, are considered for our analysis to determine the optimal window size, alternating between epochs of ∼2, 3, 4 and 5 min (i.e. averaging of 47, 71, 93 and 117 FIDs in one single spectrum) within the same wake or sleep state, i.e. wake, N1, N2, or N3. If any of these stages lasted shorter than the averaging window, the data was not included.

### Quality control of epochs

Before spectra were aligned and analyzed using LCModel, all transient of each epoch (with the final number of transients as determined in section “Trading spectral quality for temporal resolution”) were visually inspected for i) consistency of water peak frequency (wpf) location, ii) width of wpf and iii) contrast of wpf to the surrounding ppm-space. Based on these criteria, each epoch was assigned a score ranging from 1 to 5, with 1 being of high quality (i.e. wpf location consistent across the epoch, narrow wpf, and sharp contrast between wpf and surrounding ppm-area) and 5 of extreme low quality; see Supplementary Figure 1 for an example of both incidences in the same recording of one participant. Epochs with a score of ≥4 were excluded from analysis, resulting in inclusion of 95.4% (mean value; standard deviation: ±6.5%; across all participants’ epochs in the analysis. The rate of exclusion was similar between data acquired in Experiment 1 [Sleep] and Experiment 2 [Wake].

**Supplementary Figure 1.**
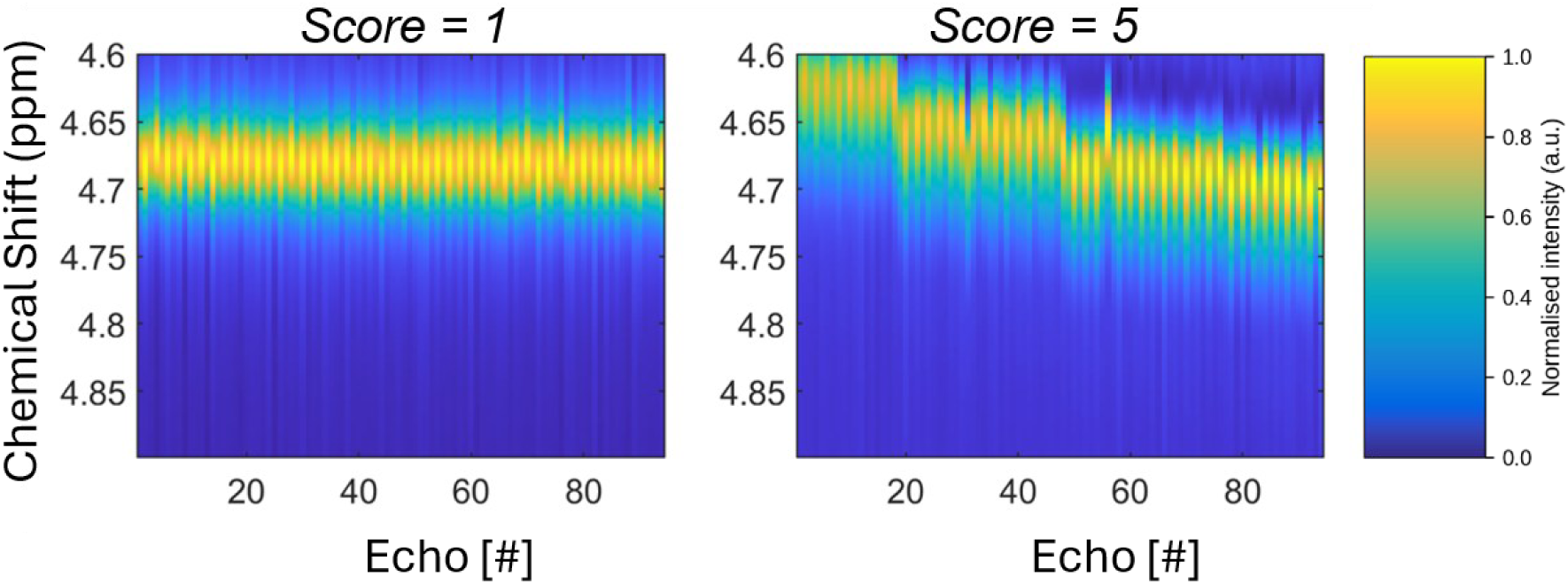
High and poor quality epochs. Example of two epochs (i.e., 93 Echo’s combined, also referred to as transients), one showing high (left) and one showing low (right) quality. The high-quality epoch is characterized by a sharp and consistent water peak frequency (wpf, around ∼4.68) on the y-axis, a narrow peak and sharp contrast with signals outside of the expected wpf. The low-quality epoch is characterized by abrupt shifts of the wpf, a broad peak and diminished contrast with signal surrounding the supposed chemical shift of water. Both epochs come from the same participant.

### Merging of EEG with MRS data

The EEG-data from the sleep condition was aligned in time with the MRS data using Brain Vision software. Specifically, the MR scanner sends a trigger upon specific events which allows to re-align the EEG signal to the FIDs created by the MR scanner. After assigning sleep-wake states to the EEG data, which was performed for the previous study ^19^, data is exported as a .txt file indicating for each FID the corresponding vigilance state.

### Statistical analyses

Results are reported throughout the manuscript as mean values ± standard deviation, unless otherwise mentioned. Data are plotted in GraphPad, Prism 10.3.1. Statistical analyses were conducted in Prism 10.3.1 and R, and are specified in the figure legends or text. To test for statistical differences in our time course data with a repeated measures ANOVA-like approach, we used mixed linear modelling (package lme4) allowing for missing values and nested designs. Details of these mixed linear models, including input parameters and outcomes, are reported in the supplementary data.

## Results

### 1. Predicting brain temperature using MC in sheep head

Temperature decreased from close-to/nearly physiological brain temperature values shortly after decapitation by ∼5°C after 3 hours (Figure 2A), and the rate of the temperature decrease was similar across samples over the course of scanning (Figure 2A, see also slope in figure legend). The Δ chemical shift increased over the course of the experiment as the brain cooled down (Figure 2B), meaning that the chemical shift of water had increased and moved further away from NAA. Hence, there was a strong correlation between temperature and Δ chemical shift with an R^2^>0.96 for all three samples (Figure 2C). In conclusion, our MRS sequence and data processing facilitates mapping of a proxy of brain temperature, showing a strong correlation with probe measurements and allowing reliable tracking of temperature changes within a sample over time.

**Figure 2.**
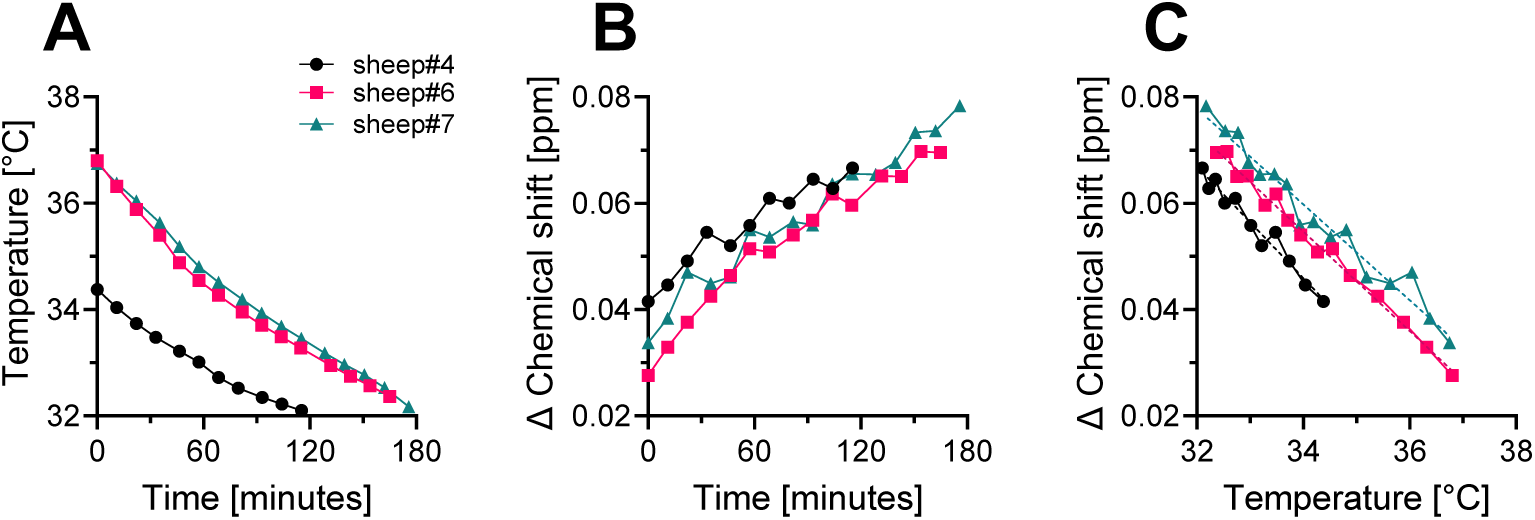
Using Δ chemical shift to predict changes in temperature. A) Brain temperature decrease in severed sheep heads across the experiment (Δ°C/minute, sheep 4: -0.020 sheep 6: -0.026; sheep 7: -0.025). B) increase in Δ chemical shift across the experiments (Δ ppb/minute, sheep 4: 0.21; sheep 6: 0.24; sheep 7: 0.23). C) Strong association between temperature measured with optical fiber and Δ chemical shift; all R^2^>0.96; dotted lines represent the correlation.

### 2. Trading spectral quality for temporal resolution

Next, we investigated how spectral averaging size (approximately 2, 3, 4, and 5 min reconstructed averages) affected linewidth, SNR, and CRLB in the in vivo MRS data. Quality metrics were averaged within each participant and subsequently summarized across participants as mean ± SD. The linewidth is not affected by the different averaging sizes (Figure 3A), but the SNR increases significantly with an increasing number of FIDs added to the epoch (SNR 2min: 12.0±1.9, 3min: 14.4±2.3, 4min: 16.5±2.7, 5min: 18.1±3.0) indicating improved signal quality. The number of participants with a mean SNR below our threshold of 12, decreases with increased averaging size (2 min: 7/12 participants; 3 min: 2 participants; 4 min: 1 participant; 5 min: 0 participants; Figure 3B). SNR gains were 20% (2→3 min), 15% (3→4 min), and 9% (4→5 min), indicating diminishing returns of the improvement curve beyond 4 min as expected according to expected SNR gains when including more datapoints. CRLB, which reflects goodness-of-fit and shows lower values with improved fitting, is decreasing when increasing the averaging window size, and reaching a significant decrease at 4 min relative to 2 min of averaging (CRLB 2 min: 9.7±7.4, 3min: 6.7±1.3, 4min: 6.1±1.2, 5min: 5.7±1.1; Figure 3C).

**Figure 3.**
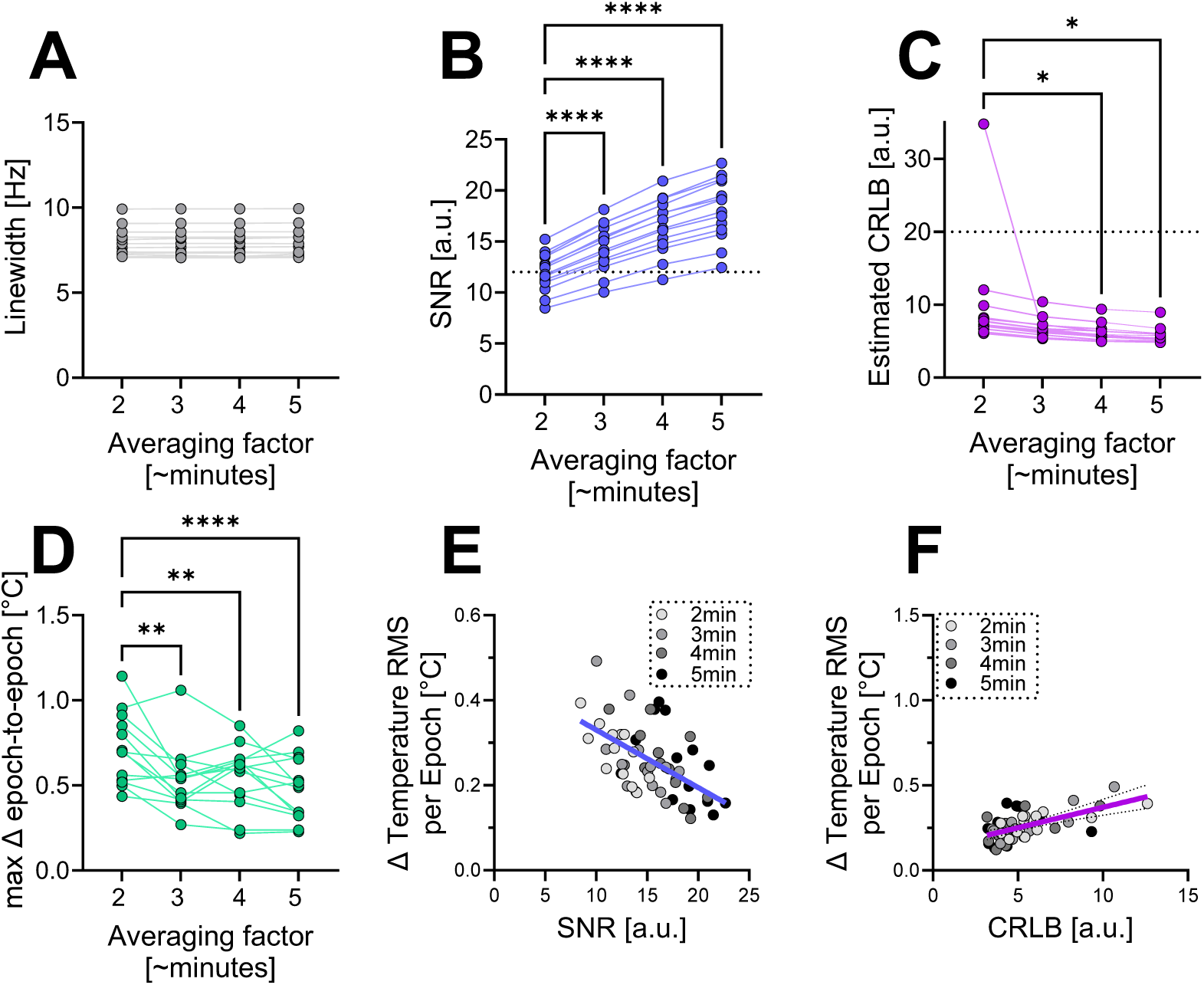
Relationship between averaging window size and. **A)** linewidth of water; **B)** signal-to-noise ratio (SNR), RM 1-W ANOVA, F(3,39)=412.5, P<0.0001, post-hoc Sídák’s MCT; **C)** estimated Cramer-Rao Lower Bound (CRLB), 1-W ANOVA, F(3,39)=3.3, P=0.0313; **D)** maximum absolute difference in estimate of brain temperature detected from epoch to epoch per participant, mixed-effects model, F(3,38)=9.65, P<0.0001, post-hoc Šídák’s MCT). **E)** Linear regression between Δ Temperature root mean square (RMS) and SNR F(1,53)=24.1, P<0.0001, R^2^=0.31 (); **F)** Linear regression between Δ Temperature RMS and CRLB (F(1,53)=27.7, P<0.0001, R^2^=0.34). Post-hoc, *:P<0.05; **:P<0.01 ****: P<0.0001.

We then inspected how variability in temperature estimates correlates with changes in spectral quality. To this end, we first quantified the maximum epoch-to-epoch variation in estimates of brain temperature per recording per averaging windows, and found that longer averaging windows are associated with smaller maximum epoch-to-epoch temperature differences (mean ± SD across participants: 2 min, 1.02 ± 1.17°C; 3 min, 0.54 ± 0.19°C; 4 min, 0.55 ± 0.18°C; 5 min, 0.48 ± 0.18°C; Figure 3-D). Δ temperature RMS, which quantifies the variability of temperature estimates across all epochs, correlates negatively with SNR (Figure 3E). Likewise, higher CRLB values, reflecting lower spectral quality, were associated with increased Δ temperature RMS. Notably, the association between Δ temperature RMS and CRLB weakened with increasing averaging windows and was no longer significant at an averaging window of 5 min.

Based on these results, we decided to use an averaging window of 4 min: this is at the point where a gain in SNR is not improving in a meaningful way when including more transients, and only one recording (out of 14) is not meeting the SNR recommended threshold of >12 at this point. Moreover, CRLB is becoming significantly smaller compared to the 2 min averaging data at 4 min averaging. Last, the correlation between variation in temperature estimates that is unlikely to be physiological, and quality measures of fitting has weakened at this point. For these reasons, an averaging window of 4 min appears a good compromise between temporal resolution and data quality.

#### From static to moving average data

To increase our temporal resolution without sacrificing spectral resolution, we assessed the possibility of using a moving average across 4-minute windows, with a window advance every 1 minute, independent of state. This method will capture more faithfully the expected transient changes in brain temperature that could otherwise be averaged away. We compared our read-outs of quality control (average SNR and estimated CRLB) from the static condition at 4 min resolution (as in Figure 3) with the moving average (MA) and did not observe any meaningful differences (SNR static: 16.5±2.7, MA: 16.6±2.7; CRLB static: 6.0±1.2, MA: 6.1±1.2). Hence, our results do not indicate that using a moving average causes loss of data quality, and we therefore decided to use a moving average with a window of 4 min and step-size of 1 minute for our further analysis. To achieve this, we merged 93 FIDs, reflecting 4 min, in one epoch, every minute. This combination of FIDs was used to estimate the chemical shift of water and NAA using LC-model.

### 3. Description of sleep-wake states in the scanner

Sleeping in the MRI scanner requires participants to be immobile and to fall asleep while the switching of magnetic field gradients creates loud noises. After launching the sequence, sleep onset latency, defined as spending at least 6 min in N2 sleep, was variable (30±22.5min; Figure 4A), and stages N4 and REM sleep were not reached. Sleep offset, defined as awakening after which no continuous episode of 10 min of N2 or N3 sleep happened, occurred on average one hour after sleep onset (average time spent “asleep”: 54.9±26.1min, Figure 4A–B). Only one participant woke up during the sleep episode (reflected by a short wake after sleep onset Figure 4C) and went back to sleep, and two participants spent time in N1. The majority of time during sleep was spent in N2 (all participants) and N3 (8 out of 9 participants) (Figure 4C–E).

**Figure 4.**
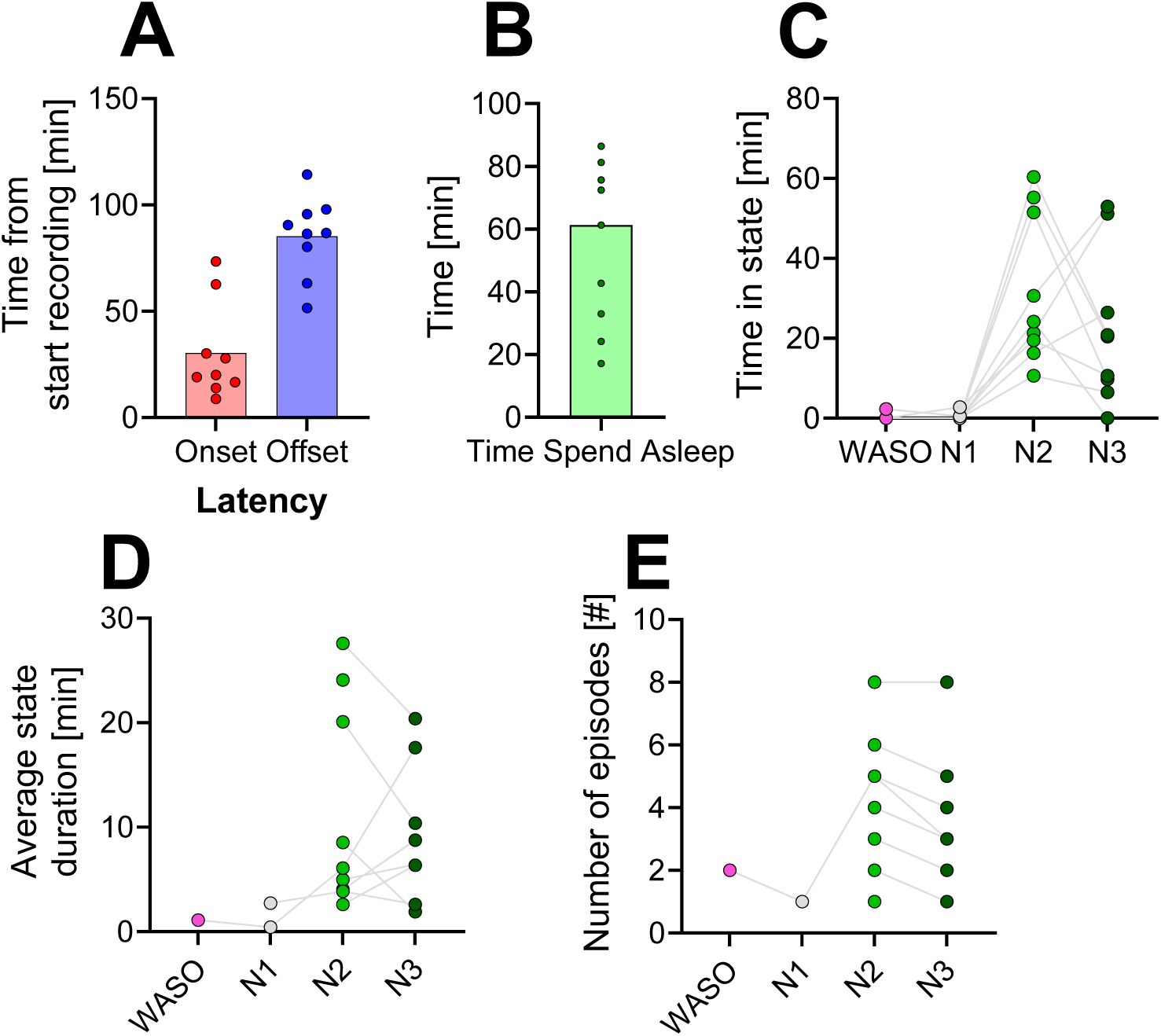
Sleep patterns in the MRI scanner (Experiment 1). **A)** Time of sleep onset and sleep offset relative to MRS recording start; **B)** Sleep episode duration (time between onset and offset) **C)** Time spent per state after sleep onset. **D)** Mean duration of time spent in each state between sleep onset and offset, **E)** Number of episodes in each state between onset and offset. Each datapoint represents one participant (n = 9). Missing values indicate that a state was not detected between sleep onset and offset in a participant. WASO = wakefulness after sleep onset, N1 – 3: NREM sleep stages N1 – N3.

### 4. Estimates of brain temperature prior to sleep

Core body temperature varies between individuals (>2° ^25^) but is relatively stable within individuals (SD ∼0.3°C; ^26^). A similar pattern was found in brain temperature measured by MRS (see for between variability: ^18^ and for within stability ^27^). Hence, we evaluated whether there was within-participant stability of brain temperature measurements across our experiments and compared estimates of brain temperature during the waking period preceding sleep onset in Experiment 1 (sleep) and the matching time window of Experiment 2 (control – wake only). We observed a significant correlation (Figure 5A, R^2^=0.29) and mean estimates of brain temperature during this initial wakefulness period did not differ significantly between experiments when incorporating all participants [Wake] (Figure 5-B, Experiment 1: 38.5±0.4°C, Experiment 2: 38.4±0.4°C; unpaired t-test, t(12)=0.58, P=0.57). However, average estimates of brain temperature were lower in all participants participating in Experiment 2 [Wake] compared to Experiment 1 [Sleep] (Figure 5B) by ∼0.25°C lower in Experiment 2 (mixed linear model, (F(1,272)=111.43, P<0.0001, details in supplementary statistics table). These measurements were taken in the participants whilst still being awake in both experiments and differences in sleep-wake states can thus not underlie this systematic variation. Importantly, this observation underscores the significance of inspecting relative changes in estimates of brain temperature within an experiment. Therefore, we next inspected relative changes in estimates of brain temperature correlating with changes in sleep-wake states.

**Figure 5.**
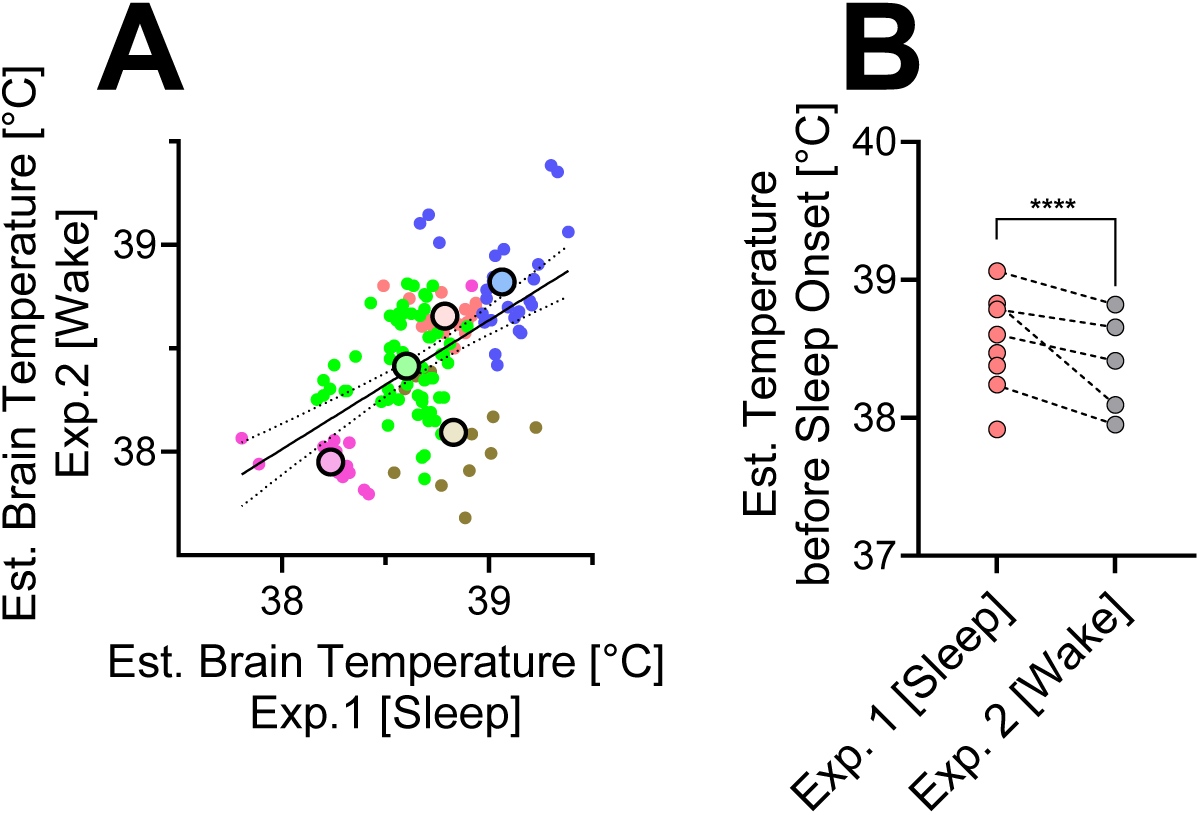
Estimates of brain temperature between experiments within participants. **A)** Brain temperature estimated per minute in the first experiment correlates with time-matched brain temperature estimates in the second experiment; each datapoint represents one epoch; epochs coming from the same participants are color-coded with the same color. Data is plotted for the participants present in both Experiment 1 and 2 (n=5). Large dots represent the mean measured temperature for one participant. Linear regression, F(1,137)=56.1, P<0.0001, R^2^=0.29 **B)** Estimates of average brain temperature prior to sleep onset in Experiment 1 and time-matched control from Experiment 2. ****, P<0.0001 (mixed linear model).

### 5. Sleep-wake evoked changes in estimates of brain temperature

To assess sleep-evoked changes in estimates of brain temperature, we first visualized estimates of brain temperature together with sleep-wake states (Supplementary Figure 2). The figure shows both variability in sleep (state) specific time (as eluded to in Figure 3), as well as variability in estimates of brain temperature, including in the absence of sleep (Supplementary Figure 2; graphs of last 5 participants).

**Supplementary Figure 2.**
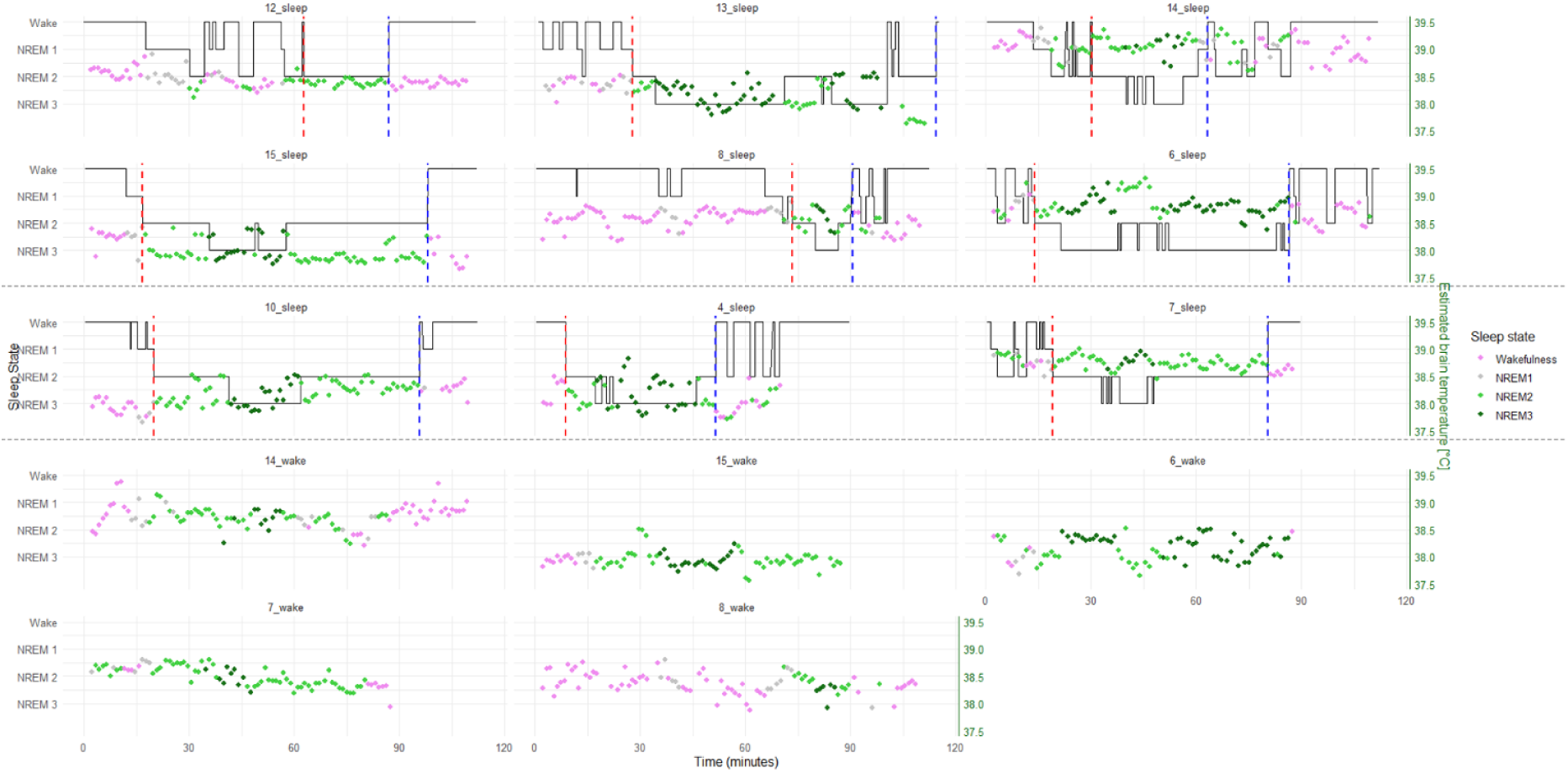
Sleep-wake state and estimates of brain temperature visualized for all participants. Participant-ID and experiment indicated on top of graph, and their sleep-wake state, visualized as hypnogram (y-axes on left hand) visualized together with estimates of brain temperature (y-axes on right hand). Estimates of brain temperature are plotted ∼ every minute, and their color also indicates the occurrence of which sleep-wake state (see legend left side figure). The same color coding has been applied to participants who stayed awake in the scanner in Experiment 2. Note that epochs with a quality score of >=4 have been excluded, explaining the missing data in e.g. participant 4_sleep towards the end of the recording. Top six graphs (first box): participants who show a decrease in brain temperature during NREM sleep 3 (dark green) compared to wakefulness (magenta) prior to sleep onset; center 3 graphs: no visual indication of a decrease in estimate of brain temperature during NREM sleep; bottom 5 graphs: brain temperature estimates measured during control Experiment 2 (Wake). Red vertical line: onset of sleep (i.e. 6 minutes of sustained NREM-2); blue line: offset of sleep.

Therefore, we focused on the temporal dynamics in estimates of brain temperature around sleep-wake state transitions. A state (wake or sleep [N1-N3]) had to be uninterrupted for at least 5 min on both sides of the transition to be included. With these requirements, we detected five transitions in Experiment 1 [Sleep], and found three matching transitions in Experiment 2 [Wake]. Across the analyzed wake-to-sleep transition period, estimates of brain temperature were approximately 0.25°C lower in Experiment 1 [Sleep] compared to the time-matched wake control group (Experiment 2) (Figure 6A, mixed linear model, F(1,154.5)=14.3, P=0.0002, Supplementary Statistics File). However, the change in temperature estimates over time did not differ significantly between the two experiments. This analysis was further limited by the variability in the duration of sleep episodes. For example, 10 min post sleep-onset, one participant woke up, and 5 min later another participant as well, reducing the number of participants contributing to the average time course over time. Moreover, only 5 of 9 participants met the criteria of having a consolidated episode of wakefulness and sleep on both sides of the transition.

**Figure 6.**
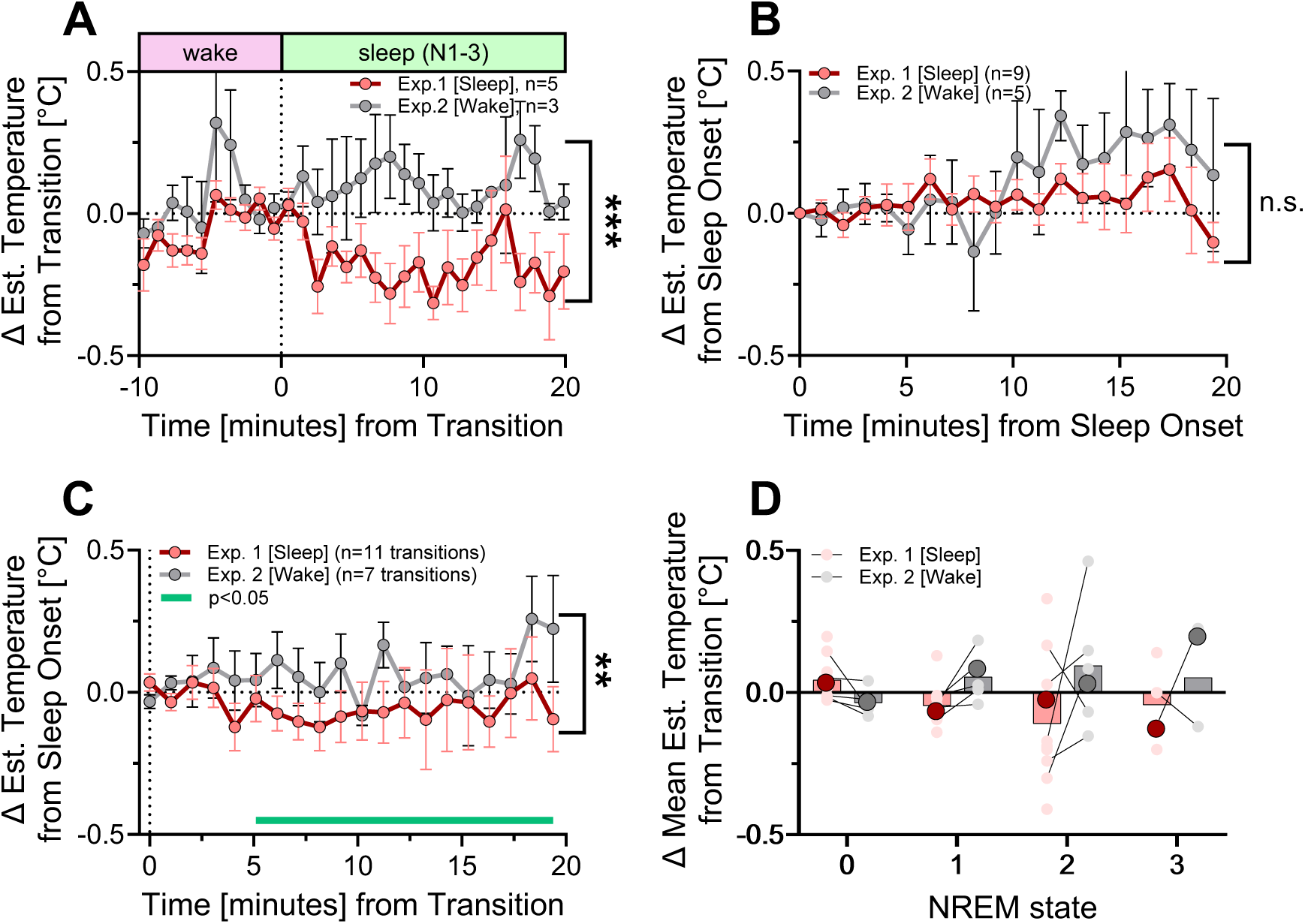
Subtle sleep-evoked decreases in estimates of brain temperature. **A)** Wake-to-sleep transitions consisting of at least 5 minutes of wakefulness before and sleep after the transition. Estimates of brain temperature differed significantly between experiments (***, P=0.0002), whereas their change over time did not differ significantly between conditions. **B)** Dynamics in estimates of brain temperature from sleep onset onwards, non-significant effect of group (P=0.61). **C)** A more inclusive analysis of estimates of brain temperature upon entering NREM sleep, including transitions into at least 5 minutes of sleep. (**, P=0.008, post-hoc defining specific times when Sleep differs significantly from Wake, P<0.05). **D)** Assessing the impact of sleep-specific states on estimates of brain temperature, based on the data in **C**. Change in temperature estimates from the last wake epoch (=0) into entering the different NREM states (1-3). Each light colored dot represents the average of one participant, whereas the large dot represents the average of the state independent of participant. Significant interaction between Experiment and state, P=0.014. All time course data are plotted as mean ± SEM. Exp = Experiment.

Therefore, we created a more inclusive analysis and focused on the changes in estimates of brain temperature from our previously defined sleep onset (Figure 6A), excluding a sustained episode of prior wakefulness as a prerequisite. Estimates of brain temperature were tracked from sleep onset and continued when the participants entered a deeper NREM sleep state (if applicable) until they exited this state (either from N3 -> N2; eight out of nine participants; or N2 -> N1; one participant). By focusing on time points that included either a stable or deepening of sleep, but not a transition into a lighter sleep state, we aimed to exclude any additional variation induced by reverse state-switching. With these criteria, we detected in our nine participants of Experiment 1 [Sleep] an average sleep duration of ∼21.8±11.9 min, ranging from 9.18 min in two participants until 39.8 min as the longest period of sleep. We matched the sleep episodes with the participant wake-matched control time from Experiment 2. Although the mean of the relative change in estimated brain temperature during sleep (Experiment 1) was below that of the wake group (Experiment 2), no significant difference in brain temperature estimates was detected between the two experiments (Figure 6B) (mixed linear model, effect of experiment, P=0.61).

We observed though that many participants slumbered into sleep and woke up again before their defined sleep onset (e.g., participant 8, 13 – see hypnograms in Supplementary Figure 2). The transition analysis in Figure 6B started from this defined sleep onset onwards and did not include NREM sleep occurring prior to the sleep onset, whilst sleep-dependent changes in the peripheral thermoregulation, i.e. heat dissipation through hands and feet, are already well on their way at this time ^3^. Hence, we expanded our transition analysis and included all transitions from wake (minimal one epoch, included as a reference) to sleep (minimal five epochs ∼5 min), independent of occurrence relative to N2 sleep onset.

By using these more complete criteria, we detected a significant sleep-evoked decrease in estimates of brain temperature over time (Figure 6C, mixed linear model; F(1, 248)=7.14, P=0.008 – see Supplementary Statistics file). The type of NREM sleep states varied significantly between participants, which is not accounted for in any of the analyses so far and thus might ignore subtle, state-specific evoked changes in estimates of brain temperature. Hence, we wondered whether there was a state-specific effect on brain temperature estimates and assessed this question in the data set containing the most datapoints (Figure 6C). By comparing the effect of different NREM sleep states on estimates of brain temperature, with time-matched control estimates from wakefulness (i.e. comparing Experiment1 [Sleep] vs Experiment 2 [Wake]), we found a significant interaction between the two: the deeper NREM sleep, the more pronounced the decrease in brain temperature in sleep, relative to the time-matched wake control condition (Figure 6D, mixed linear model; F(1,3)=244.9, P=0.014 – see supplementary statistics file). Taken together, these results support the existing literature in rodents and the correlative evidence in humans that during NREM sleep, brain temperature decreases.

## Discussion

We hypothesized that thalamic temperature estimates would decrease during NREM sleep compared with time-matched wakefulness. Consistent with this hypothesis, thalamic temperature estimates decreased by approximately 0.2°C upon entering NREM sleep during an evening nap, with a more pronounced decrease during deeper NREM sleep. This finding is consistent with observations in non-human mammals, in which brain temperature decreases during NREM sleep, and with the only direct human recordings obtained invasively in patients with epilepsy ^11^.

### Temperature variation across the day

Our data are in line with the results from Landolt et al. ^11^, who observed a decrease in parahippocampal gyrus temperature in epilepsy patients during sleep. Intriguingly, in this study, the decrease in brain temperature was already initiated at lights off prior to the participants entering NREM sleep which suggests that independent of sleep itself, thermoregulatory changes to initiate brain cooling occur already in the sleep preparatory phase. An important sleep-preparatory change is body posture (i.e. going into a supine position), which reduces tympanic temperature, an established proxy of brain temperature ^28^. In our experimental design, we controlled for the sleep-associated change in posture with a wake control experiment. Moreover, with this control group, we also excluded a potential time-of-day effect, which is important because the temperature of the incoming blood towards the brain, reflected by core body temperature, is one of the major determinants of brain temperature, and is under strong control of the circadian clock ^29^. Along these lines, Rzechorzek and colleagues ^18^ found a significant time-of-day effect on estimates of brain temperature using MRS, with brain temperature varying up to 0.8°C across the day. In their study, estimates of brain temperature were lowest in the evening. This could indicate that unlike in small rodents, where brain temperature variation is driven by sleep-wake state ^8,30^, in humans, brain temperature is the resultant of an interaction with circadian time and sleep-wake state. Further experiments using a forced desynchrony protocol will disentangle the contribution of these factors to brain temperature dynamics.

### Mechanisms contributing to sleep-evoked changes in brain temperature

Brain temperature variation can be roughly understood by the contribution of two factors: heat production by neural cells, and the efficiency at which this heat can be dissipated. During NREM sleep, cortical glucose metabolism is lower compared to wakefulness ^31^, in line with slower neuronal oscillations during this vigilance state and increased “OFF” periods during which glutamatergic neurons remain silent ^32^, requiring less energy and thus reducing the amount of heat that is produced. Indeed, in our data we observed a sleep-state dependent decrease in brain temperature (Figure 6D), where brain temperature was significantly more decreased when sleep was “deeper”, i.e. NREM-1 versus NREM-3. In rodents, there is limited evidence that spectral composition, reflecting neuronal activity, during NREM sleep correlates with changes in brain temperature: NREM sleep spindles can lead to a subtle but consistent increase in temperature ^9^, but on a longer time-scale, there is no correlation between hourly NREM slow-wave activity and average cortical temperature (PMID: 1858947). In our data, we cannot decompose the EEG signal into different frequency bands due to the artefacts induced by the MR scanner. However, considering new developments in EEG-acquisition paradigms that allow for cleaner EEG-data to be obtained, future experiments co-detecting EEG-based brain activity and estimates of brain temperature, can dissect at higher resolution the contribution of neuronal activity to brain temperature changes.

Next to neuronal activity, the temperature of the incoming peripheral blood, reflected by core body temperature, is the other major determinant of brain temperature. Already ∼2 hours before sleep onset, thermoregulatory changes are initiated, causing a decline in core body temperature ^33^. Specifically, heat is distributed away from the core towards the shell (i.e. extremities), and this effect becomes amplified when sleep preparatory behavior is initiated, with increased vasodilation in so-called glabrous skin areas (feet- and hand palms, forehead). This process can be sped up by promoting vasodilation by e.g. wearing socks ^5^, whereas people with impaired vasodilation suffer from sleep issues, including prolonged time to fall asleep ^4^, underscoring that sleep and thermoregulation are tightly regulated together. The cooler blood entering the brain during NREM sleep can then dissipate heat more efficiently, further enhancing cooling of the brain. To what extent either of these factors (heat production vs dissipation) contributes to the sleep-dependent changes in brain temperature is not understood and requires mathematical modelling combined with functional experiments.

### Thalamic temperature: a proxy for global brain temperature?

Brain temperature varies by location in the brain: the deeper in the brain, the warmer the brain. This phenomenon has been measured in the early experiments from Hayward and Baker in animal studies ^34,35^, and was confirmed in humans ^18^. This temperature gradient can be understood through different heat dissipation characteristics at different depths within the brain: Deeper brain structures cannot passively dissipate heat through the skull as they are surrounded by other, warm brain tissue and are therefore warmer compared to superficial structures such as the cortex. Despite these regional differences in absolute temperature, the impact of sleep on relative brain temperature across brain regions is very similar: For example, in the rat, the daily amplitude in the cortex is 1.1°C and 1.0°C in the hypothalamus ^36^. Whether in humans, the impact of sleep on different brain regions is uniform, is not known.

### Technical limitations

Our study is the first that investigated sleep-evoked changes in human brain temperature by carefully controlling for the confounding factors of time-of-day and body posture. In this section, we would like to address limitations of this study. First, the small sample size (n=9 for sleep) brings challenges as their sleep varied widely: for example, sleep onset was reached in one participant after 8 min, whereas it took another participant 73 min to reach stable NREM sleep. This inter-individual variability makes it rather challenging to draw solid conclusions on the sleep-evoked variation in human brain temperature. This limitation is difficult to overcome because sleeping in the MR environment is difficult *per se* and participants respond differently to this challenge. Second, of the nine participants who were able to sleep, only 5 managed to stay awake in the wake control condition, which reduces statistical power of the control group. Third, Experiment 1 [Sleep] and Experiment 2 [Wake] were not conducted in a randomized order, making it challenging to understand whether the consistently lower detected estimate of brain temperature in Experiment at the baseline measurement (i.e. during wakefulness prior to sleep) could be due to e.g. a seasonal effect, or a technical aberration in the MR scanner. By increasing the sample size and randomizing timing of experimental and control conditions in further experiments, these shortcomings can be overcome in future studies. Fourth, as this study was based on retrospectively analyzed MRS data that had originally been acquired for a different research purpose, voxel size and placement were predetermined by the original study protocol. The resulting thalamic voxel constitutes a technically demanding setting for MRS-based thermometry. A larger voxel placed in a cortical region would be expected to provide higher SNR within the same acquisition time. As temperature estimation relies on the precise determination of resonance frequencies, increased SNR would likely improve both measurement precision and temporal resolution. Last, motion-related effects may have contributed to measurement variability. Owing to the anatomical complexity of the thalamic region, even subtle subject movement can alter the relative tissue composition within the voxel and affect spectral quality. Such effects may be less pronounced in cortical regions with more homogeneous tissue composition, where comparable voxel displacements would be expected to have a smaller impact on spectral characteristics and, consequently, on temperature estimates.

## Conclusion

Our results point towards a subtle sleep-induced cooling during NREM sleep relative to wakefulness, and we note that the changes are small, and statistically significant. Our results show the feasibility of using the MRS technique in studying sleep-evoked changes in estimates of brain temperature. Future studies would benefit from a larger sample size and a full night of sleep to test whether nocturnal sleep cools the brain, and at which magnitude.

## Data availability

The data underlying the Figure 2 to Figure 6 will be made available .csv files. The R-script used to analyze the data will also be made available. Upon request, other data can also be made available.

## Acknowledgements

We would like to thank: Professor Gilles Vandewalle for insightful discussions on the data. We would like to thank Professor Michael Thali for providing the opportunity to visit the of Forensic Medicine in Zürich at University of Zürich.

## Statistics output from mixed linear models

**Figure 5A-B.**
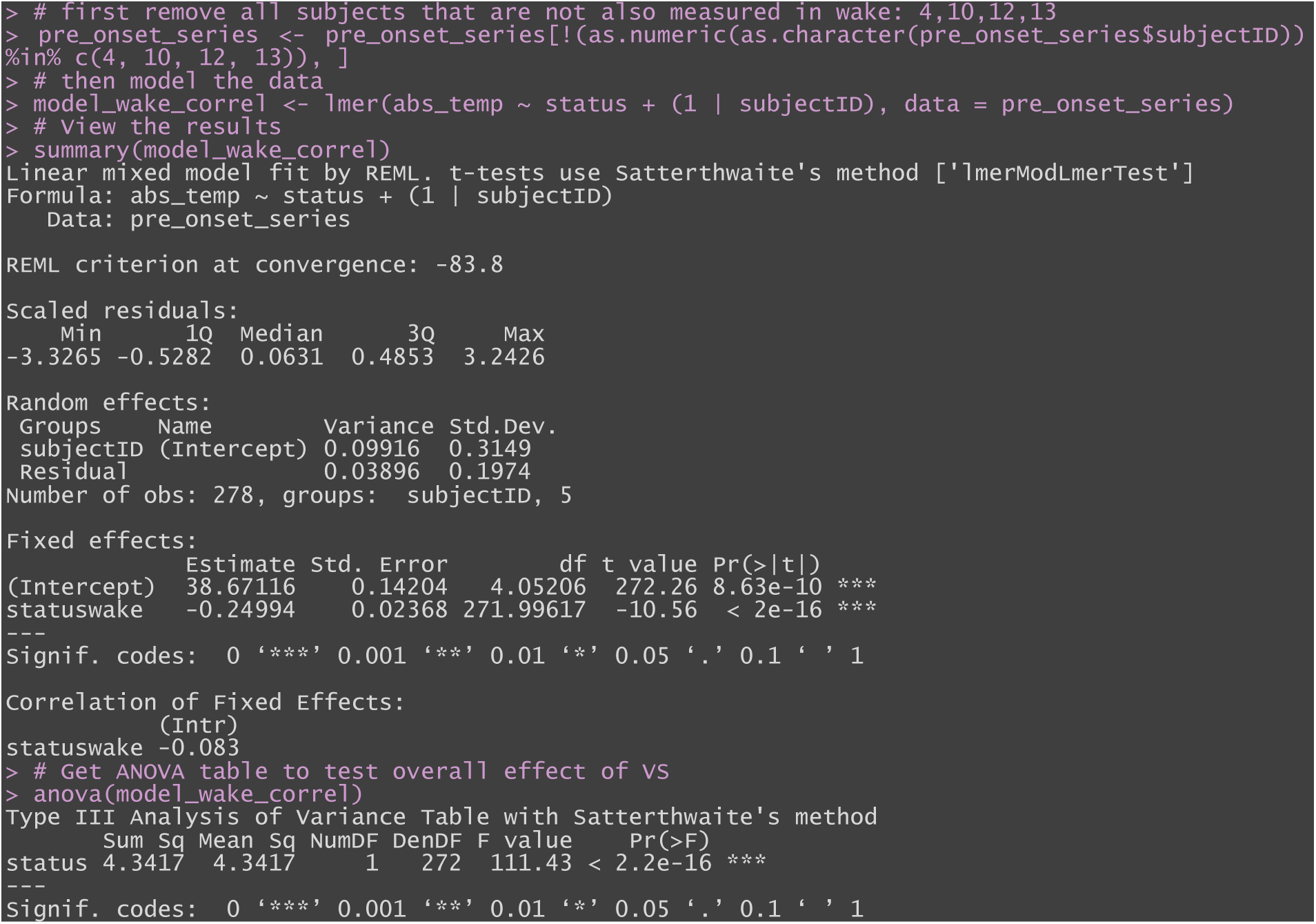
comparing estimates of brain temperature, epoch-by-epoch, prior to sleep onset in Experiment 1 vs Experiment 2 (“status”)

**Figure 6-A.**
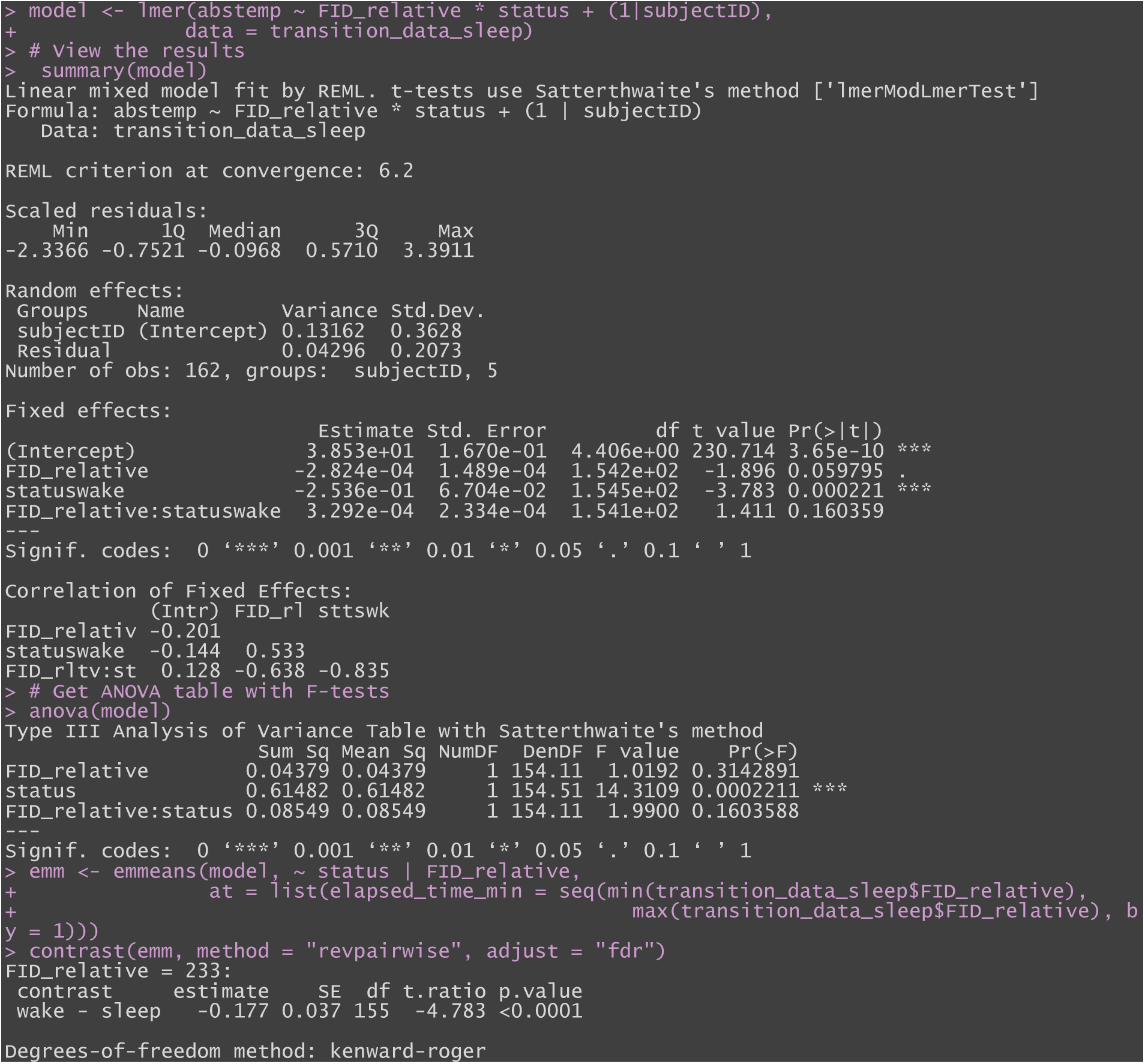
wake to sleep transition, assessing whether there is a differences relative temperature upon entering sleep (or wake-control, Exp2). Factor Experiment (i.e. Experiment 1 [Sleep] or Experiment 2 [Wake]), represented by “status”.

**Figure 6-B.**
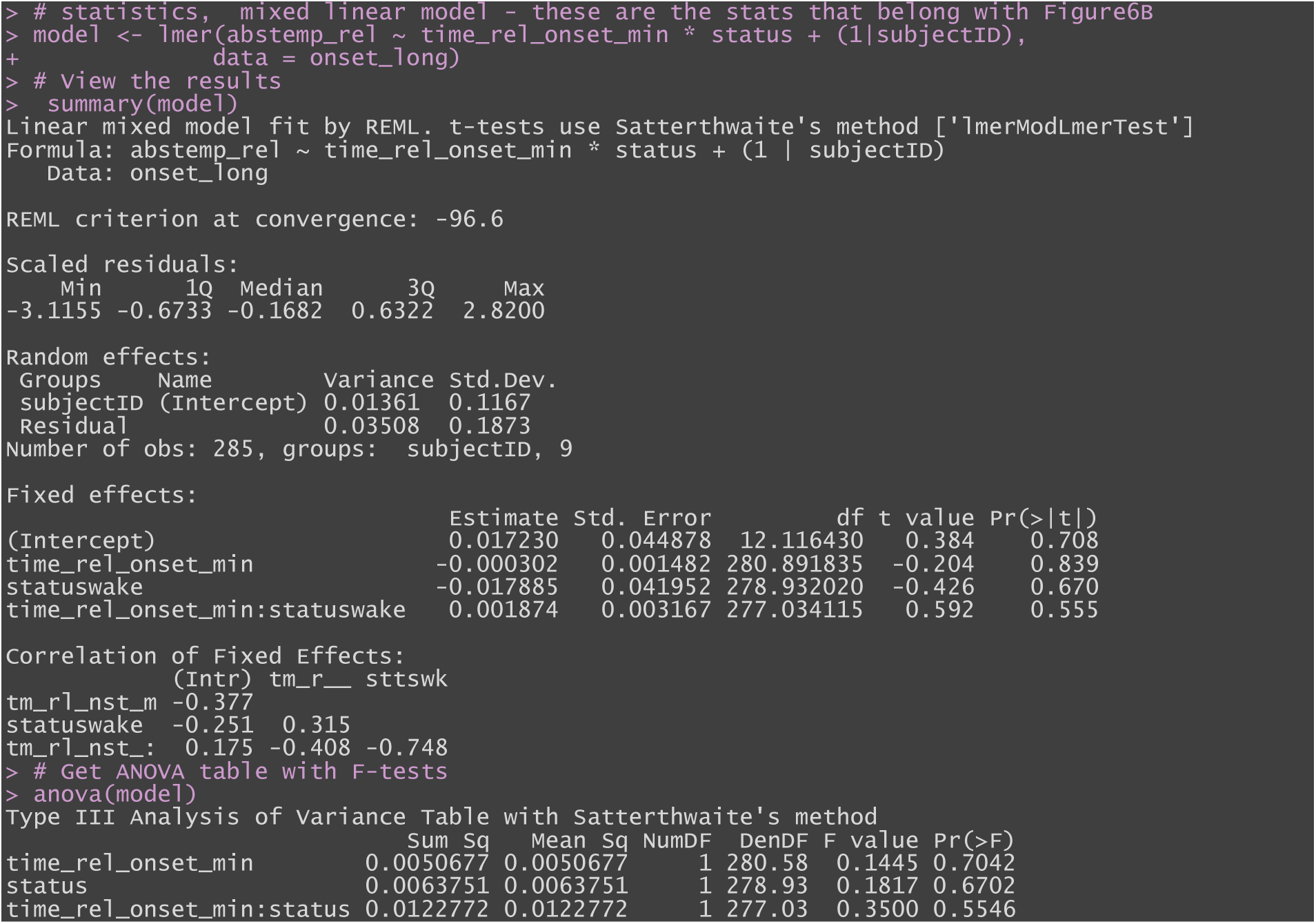
changes in relative temperature from sleep onset on. Factor Experiment (i.e. Experiment 1 [Sleep] or Experiment 2 [Wake]), represented by “status”.

**Figure 6-C.**
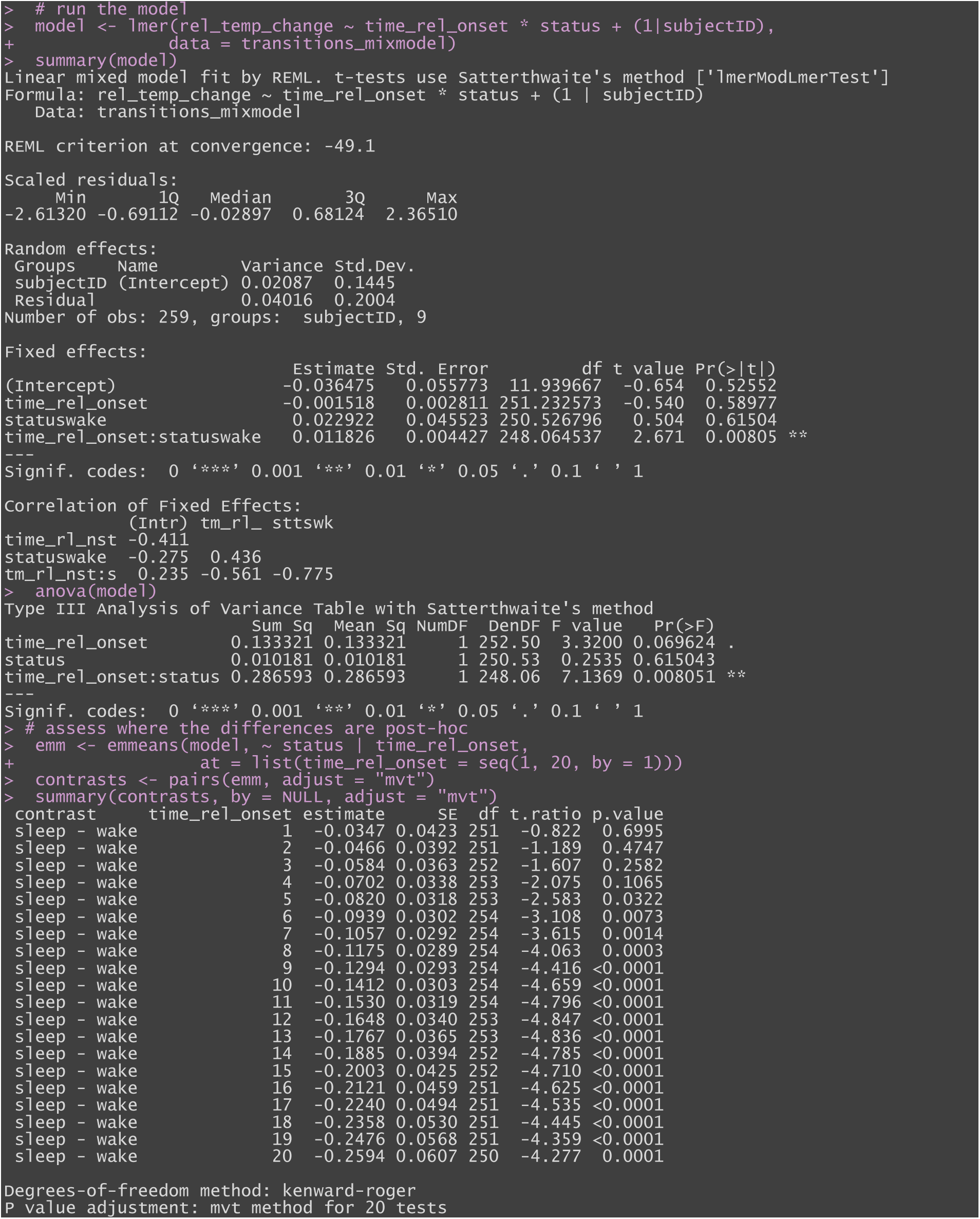
Entering sleep without the prerequisite of previous sustained wakefulness occurring any time during the recording. Sleep should last at least 5 minutes. Factor Experiment (i.e. Experiment 1 [Sleep] or Experiment 2 [Wake]), represented by “status”.

**Figure 6-D.**
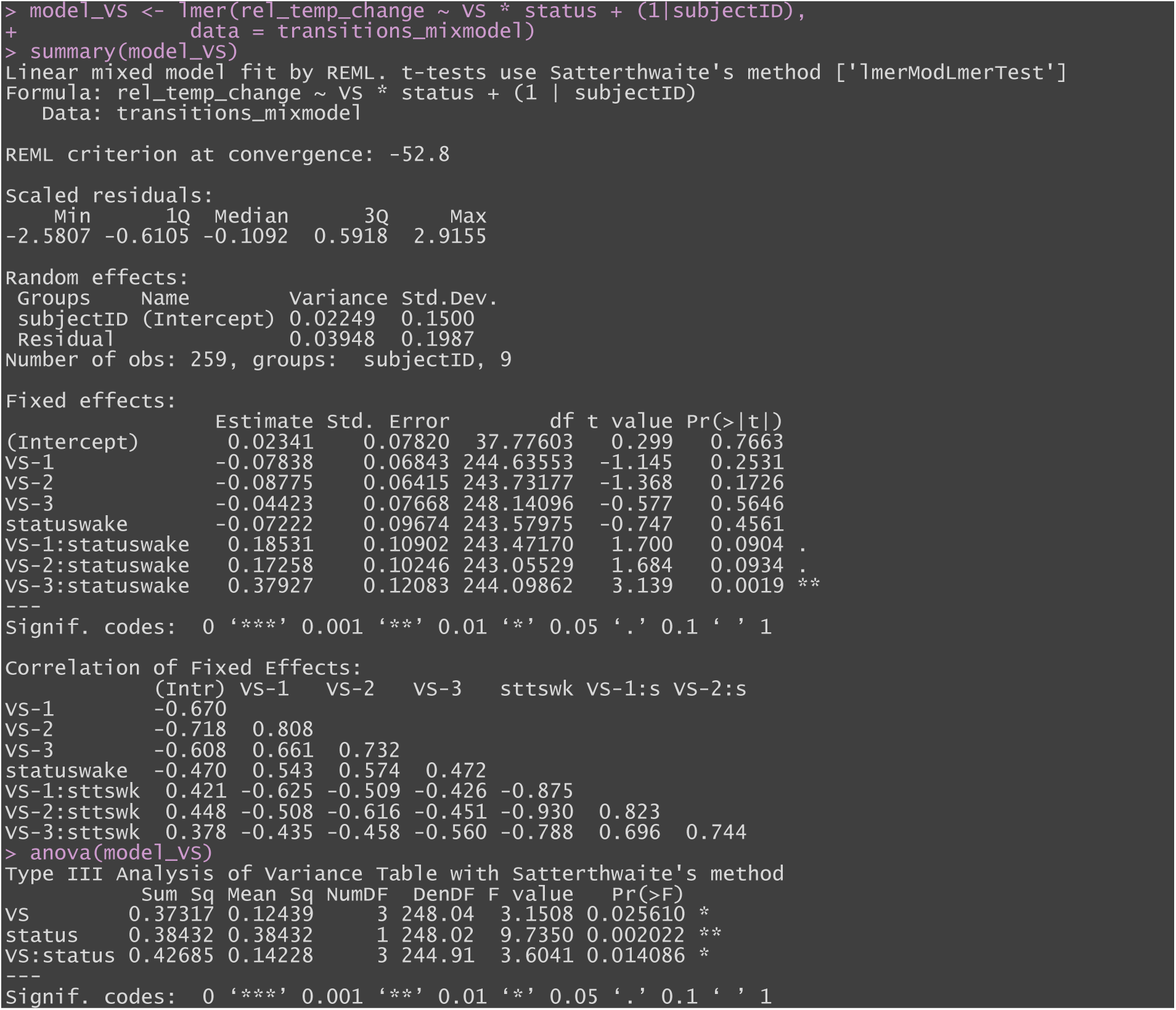
– experiment with NREM state (i.e. VS) interaction effect modelled, based on the transitions selected from Figure 6-C

## References

1. Murphy PJ, Campbell SS. Nighttime drop in body temperature: a physiological trigger for sleep onset? Sleep. 1997;20(7):505–511. doi:10.1093/sleep/20.7.505

2. Zulley J, Wever R, Aschoff J. The dependence of onset and duration of sleep on th circadian rhythm of rectal temperature. Pflugers Arch. 1981;391(4):314–318. doi:10.1007/BF00581514

3. Kräuchi K, Cajochen C, Werth E, Wirz-Justice A. Functional link between distal vasodilation and sleep-onset latency? Am J Physiol Regul Integr Comp Physiol. 2000;278(3):R741–748. doi:10.1152/ajpregu.2000.278.3.R741

4. Pache M, Kräuchi K, Cajochen C, et al. Cold feet and prolonged sleep-onset latency in vasospastic syndrome. Lancet. 2001;358(9276):125–126. doi:10.1016/S0140-6736(01)05344-2

5. Kräuchi K, Cajochen C, Werth E, Wirz-Justice A. Warm feet promote the rapid onset of sleep. Nature. 1999;401(6748):36–37. doi:10.1038/43366

6. Monroe LJ. Psychological and physiological differences between good and poor sleepers. J Abnorm Psychol. 1967;72(3):255–264. doi:10.1037/h0024563

7. Hoekstra MM, Emmenegger Y, Hubbard J, Franken P. Cold-inducible RNA-binding protein (CIRBP) adjusts clock-gene expression and REM-sleep recovery following sleep deprivation. Elife. 2019;8:e43400. doi:10.7554/eLife.43400

8. Sela Y, Hoekstra MM, Franken P. Sub-minute prediction of brain temperature based on sleep-wake state in the mouse. Elife. 2021;10:e62073. doi:10.7554/eLife.62073

9. Lazopulo A, Emmenegger Y, Đukanović N, Hoekstra MMB, Franken P. Brain temperature as proxy for brain state and oscillatory activity in the mouse. Sci Rep. 2025;15(1):37301. doi:10.1038/s41598-025-21175-3

10. Petersen PC, Vöröslakos M, Buzsáki G. Brain temperature affects quantitative features of hippocampal sharp wave ripples. J Neurophysiol. 2022;127(5):1417–1425. doi:10.1152/jn.00047.2022

11. Landolt HP, Moser S, Wieser HG, Borbély AA, Dijk DJ. Intracranial temperature across 24-hour sleep-wake cycles in humans. Neuroreport. 1995;6(6):913–917. doi:10.1097/00001756-199504190-00022

12. Cady EB, D’Souza PC, Penrice J, Lorek A. The estimation of local brain temperature by in vivo 1H magnetic resonance spectroscopy. Magn Reson Med. 1995;33(6):862–867. doi:10.1002/mrm.1910330620

13. Corbett R, Laptook A, Weatherall P. Noninvasive measurements of human brain temperature using volume-localized proton magnetic resonance spectroscopy. J Cereb Blood Flow Metab. 1997;17(4):363–369. doi:10.1097/00004647-199704000-00001

14. Kuroda K. Non-invasive MR thermography using the water proton chemical shift. Int J Hyperthermia. 2005;21(6):547–560. doi:10.1080/02656730500204495

15. Zoelch N, Heimer J, Richter H, et al. In situ temperature determination using magnetic resonance spectroscopy thermometry for noninvasive postmortem examinations. NMR in Biomedicine. 2024;37(10):e5171. doi:10.1002/nbm.5171

16. Zhu M, Bashir A, Ackerman JJ, Yablonskiy DA. Improved calibration technique for in vivo proton MRS thermometry for brain temperature measurement. Magn Reson Med. 2008;60(3):536–541. doi:10.1002/mrm.21699

17. Dehkharghani S, Qiu D. MR Thermometry in Cerebrovascular Disease: Physiologic Basis, Hemodynamic Dependence, and a New Frontier in Stroke Imaging. AJNR Am J Neuroradiol. 2020;41(4):555–565. doi:10.3174/ajnr.A6455

18. Rzechorzek NM, Thrippleton MJ, Chappell FM, et al. A daily temperature rhythm in the human brain predicts survival after brain injury. Brain. 2022;145(6):2031–2048. doi:10.1093/brain/awab466

19. Lehmann M, Hock A, Zoelch N, Landolt HP, Seifritz E. Dynamic Metabolic Changes in the Human Thalamus at the Transition From Waking to Sleep - Insights From Simultaneous Functional MR Spectroscopy and Polysomnography. Front Neurosci. 2019;13:1158. doi:10.3389/fnins.2019.01158

20. Gent TC, Bandarabadi M, Herrera CG, Adamantidis AR. Thalamic dual control of sleep and wakefulness. Nat Neurosci. 2018;21(7):974–984. doi:10.1038/s41593-018-0164-7

21. Allen PJ, Polizzi G, Krakow K, Fish DR, Lemieux L. Identification of EEG Events in the MR Scanner: The Problem of Pulse Artifact and a Method for Its Subtraction. NeuroImage. 1998;8(3):229–239. doi:10.1006/nimg.1998.0361

22. Silber MH, Ancoli-Israel S, Bonnet MH, et al. The visual scoring of sleep in adults. J Clin Sleep Med. 2007;3(2):121–131.

23. Hock A, MacMillan EL, Fuchs A, et al. Non-water-suppressed proton MR spectroscopy improves spectral quality in the human spinal cord. Magn Reson Med. 2013;69(5):1253–1260. doi:10.1002/mrm.24387

24. Landheer K, Juchem C. Are Cramér-Rao lower bounds an accurate estimate for standard deviations in in vivo magnetic resonance spectroscopy? NMR Biomed. 2021;34(7):e4521. doi:10.1002/nbm.4521

25. Obermeyer Z, Samra JK, Mullainathan S. Individual differences in normal body temperature: longitudinal big data analysis of patient records. BMJ. 2017;359:j5468. doi:10.1136/bmj.j5468

26. Diamond A, Lye CT, Prasad D, Abbott D. One size does not fit all: Assuming the same normal body temperature for everyone is not justified. PLoS One. 2021;16(2):e0245257. doi:10.1371/journal.pone.0245257

27. Sharma AA, Nenert R, Mueller C, Maudsley AA, Younger JW, Szaflarski JP. Repeatability and Reproducibility of in-vivo Brain Temperature Measurements. Front Hum Neurosci. 2020;14:598435. doi:10.3389/fnhum.2020.598435

28. Aizawa S, Cabanac M. The influence of temporary semi-supine and supine postures on temperature regulation in humans. Journal of Thermal Biology. 2002;27(2):109–114. doi:10.1016/S0306-4565(01)00023-7

29. Czeisler CA, Gooley JJ. Sleep and circadian rhythms in humans. Cold Spring Harb Symp Quant Biol. 2007;72:579–597. doi:10.1101/sqb.2007.72.064

30. Franken P, Tobler I, Borbély AA. Sleep and waking have a major effect on the 24-hr rhythm of cortical temperature in the rat. J Biol Rhythms. 1992;7(4):341–352. doi:10.1177/074873049200700407

31. Kennedy C, Gillin JC, Mendelson W, et al. Local cerebral glucose utilization in non-rapid eye movement sleep. Nature. 1982;297(5864):325–327. doi:10.1038/297325a0

32. Steriade M. The corticothalamic system in sleep. Front Biosci. 2003;8:d878–899. doi:10.2741/1043

33. Kräuchi K. The human sleep-wake cycle reconsidered from a thermoregulatory point of view. Physiol Behav. 2007;90(2-3):236–245. doi:10.1016/j.physbeh.2006.09.005

34. Hayward JN, Baker MA. Role of cerebral arterial blood in the regulation of brain temperature in the monkey. Am J Physiol. 1968;215(2):389–403. doi:10.1152/ajplegacy.1968.215.2.389

35. Hayward JN, Baker MA. A comparative study of the role of the cerebral arterial blood in the regulation of brain temperature in five mammals. Brain Research. 1969;16(2):417–440. doi:10.1016/0006-8993(69)90236-4

36. Gao BO, Franken P, Tobler I, Borbély AA. Effect of elevated ambient temperature on sleep, EEG spectra, and brain temperature in the rat. Am J Physiol. 1995;268(6 Pt 2):R1365–1373. doi:10.1152/ajpregu.1995.268.6.R1365

